# Compression-Assisted Arthrocentesis of the Knee as a Quality Improvement Intervention

**DOI:** 10.1101/395376

**Authors:** James F. Bennett, Wilmer L. Sibbitt, Philip A. Band, Sabeen Yaqub, N. Suzanne Emil, Monthida Fangtham, Roderick A. Fields, William A. Hayward, Selma D. Kettwich, Arthur D. Bankhurst

## Abstract

**Objective:** The present study reports the introduction of mechanical compression of the knee for arthrocentesis as quality improvement intervention in a procedure clinic.

**Methods:** 430 consecutive symptomatic osteoarthritic knees underwent arthrocentesis followed by corticosteroid injection (1mg/kg of triamcinolone acetonide). The first 215 consecutive knees underwent conventional arthrocentesis and injection; the quality intervention of a mechanical compression brace was introduced, and the next 215 consecutive knees underwent mechanical compression-assisted arthrocentesis follow by injection. Pain scores, arthrocentesis success, fluid yield, time-to-next-intervention, injections/year, and medical costs were measured.

**Results:** No serious adverse events occurred in 430 subjects. Diagnostic synovial fluid (≥2 ml) was obtained in 9.3% (20/215) without compression and 40.9% (88/215) with compression (p=0.00001, z for 95% CI= 1.96, Pierson). Mechanical compression was associated with a 231% increase in mean arthrocentesis volume: compression 5.3±11.2 ml, conventional 1.6±6.4 ml (CI of difference 2.0 <3.7< 5.4; p=0.00001). Time-to-next-intervention after compression-assisted arthrocentesis was longer: 6.9±3.5 months compared to conventional: 5.1±2.7 months (p<0.00001, 95% CI of difference 1.2 <1.8< 2.3). Mechanical compression was associated with a reduction in the number of corticosteroid injections administered per year: mechanical compression: 1.7±0.9 injections/year; conventional: 2.4±0.5 injections/year (p<0.00001, 95% CI of difference −0.83 < −0.70< −0.56). Mechanical compression did not increase overall yearly costs associated with management of the symptomatic knee (mechanical compression: $293.30/year/knee, conventional: $373.29/year/knee) (p<0.0001, 95% CI of difference 47 <80< 112).

**Conclusions:** Routine mechanical compression of the knee for arthrocentesis and injection is an effective bioengineering quality improvement intervention in a procedure clinic.

## INTRODUCTION

After non-pharmacologic interventions and topical and oral medications, complete arthrocentesis followed intraarticular injection of corticosteroids is often recommended for symptomatic flares and short-term relief of pain of arthritis of the knee [1-5]. Klocke and colleagues have recently reported that corticosteroid injection of the osteoarthritic knee may actually decrease cartilage degradation in the short-term as measured by biomarkers, providing indirect support for the above recommendations [1-6]. Recently, Meehan et al, Bhavsar et al, and Yaqub et al have demonstrated that mechanical compression of the knee provides more complete arthrocentesis before injection in both the effusive and non-effusive knee and suggest that mechanical compression might be a reasonable quality improvement intervention for arthrocentesis [6-9]. In the present study we determined knee arthrocentesis and injection quality measures before and after the introduction of mechanical compression as a quality improvement intervention.

## MATERIALS AND METHODS

This Quality Improvement program was formalized in our medical center in a formal effort to improve rheumatology outcomes and services with a commitment to continuous quality improvement in outpatient rheumatology as recently described recently by Chow et al [10]. The study was approved by the institutional review board (IRB) and in compliance with the Helsinki Declaration and subsequent revisions. The sequential study design was typical of quality improvement prospective cohort study with 1) measurement of baseline quality factors in consecutive traditionally treated patients, 2) introduction of the quality intervention, and 3) re-measurement of quality factors in patients after the intervention. A total of 430 knees with grade II-III osteoarthritis were included in this study. In the primary corticosteroid arm, the first consecutive 215 knees underwent conventional arthrocentesis and injection, then the quality intervention was introduced, and the second 215 consecutive knees underwent mechanical compression-assisted arthrocentesis and injection. Inclusion criteria for the study consisted of: 1) painful symptomatic grade II-III osteoarthritis knee with the patient requesting a knee injection, 2) indications for therapeutic-diagnostic arthrocentesis and/or injection, 3) indication for corticosteroid injection, and 4) formal signed consent of the patient to undergo the procedure [11]. Prior to the procedure, the presence or absence of clinical effusion was confirmed by physical examination by palpation of the extended knee for suprapatellar bursa distention, ballottement of a floating patella while applying pressure superiorly and inferiorly, and visible-palpable fluid shift with asymmetric compression. The knee was then classified as “effusive” (swollen) or “non-effusive” (dry). Arthrocentesis prior to injection was attempted in all subjects prior to injection regardless of the presence or absence of a clinical effusion.

### Arthrocentesis and Joint Injection Technique

Chlorhexidine 2% was used for antisepsis. Knee procedures were performed using the conventional lateral approaches with the one-needle multiple syringe technique [12-20]. A 22 gauge 2 inch needle (4710007050 – 22 GX2” (0.7X50 mm), FINE-JECT, Henke Sass Wolf, Kettenstrasse 1 D-78532 Tuttlingen, Germany) on a 3 ml syringe (3 ml Luer Lok syringe, BD, 1 Becton Drive, Franklin Lakes, NJ 07417, website: http://www.bd.com) filled with 3 ml of 1% lidocaine (Xylocaine® 1%, AstraZeneca Pharmaceuticals LP, 1800 Concord Pike, P.O. Box 15437, Wilmington, DE 19850-5437) was introduced into the lateral parapatellar recess of the suprapatellar bursa and if no fluid returned, into the patellofemoral joint. The knee was then compressed or “milked” by the operator’s free hand to optimize fluid return [8,9,12-17]. If fluid return was obtained and the syringe filled with synovial fluid, the intraarticular needle was left in place, the 3 ml syringe was removed, and a syringe exchange was performed with additional 20 ml syringes until fluid return ceased. Pain scores, arthrocentesis success, and fluid yield were recorded. After the fluid return ceased, a final syringe exchange was performed, and the joint was then injected through the already-placed intraarticular needle using a 3 ml syringe with 1 mg/kg triamcinolone acetonide suspension (maximum 80 mg) (Kenalog® 40, Westwood-Squibb Pharmaceuticals, Inc (Bristol-Myers Squibb), 345 Park Ave, New York, NY 10154-0004, USA). In the quality intervention group, an elastomeric knee brace (YooSoo Adjustable Knee brace, Shenzhen Shi Hai Xun Yun Wei Co., Ltd. No.203, 69 Dong, Liyuan Xin Cun, Bantian Street, Longgang, 518000, Shenzhen, China, Amazon, https://www.amazon.com) was placed and modified so the lateral suprapatellar bursa and patellofemoral joint could expand with fluid (Figures 1 and 2]. The brace applies constant mechanical compression to the medial suprapatellar bursa and the synovial compartments of the inferior knee, collapsing these compartments, and forcing fluid to the lateral suprapatellar bursa and the patellofemoral joint where the fluid could be accessed (Figures 1 and 2). Before the brace was applied, the skin surfaces of the knee were cleaned with 2% chlorhexidine. As the brace itself was clean but non-sterile, after the compressive brace was placed, the antisepsis with 2% chlorhexidine was again applied to the skin of the operative portal overlying the lateral recess of the suprapatellar bursa, and arthrocentesis and injection were performed identically as described above.

**FIGURE 1.**
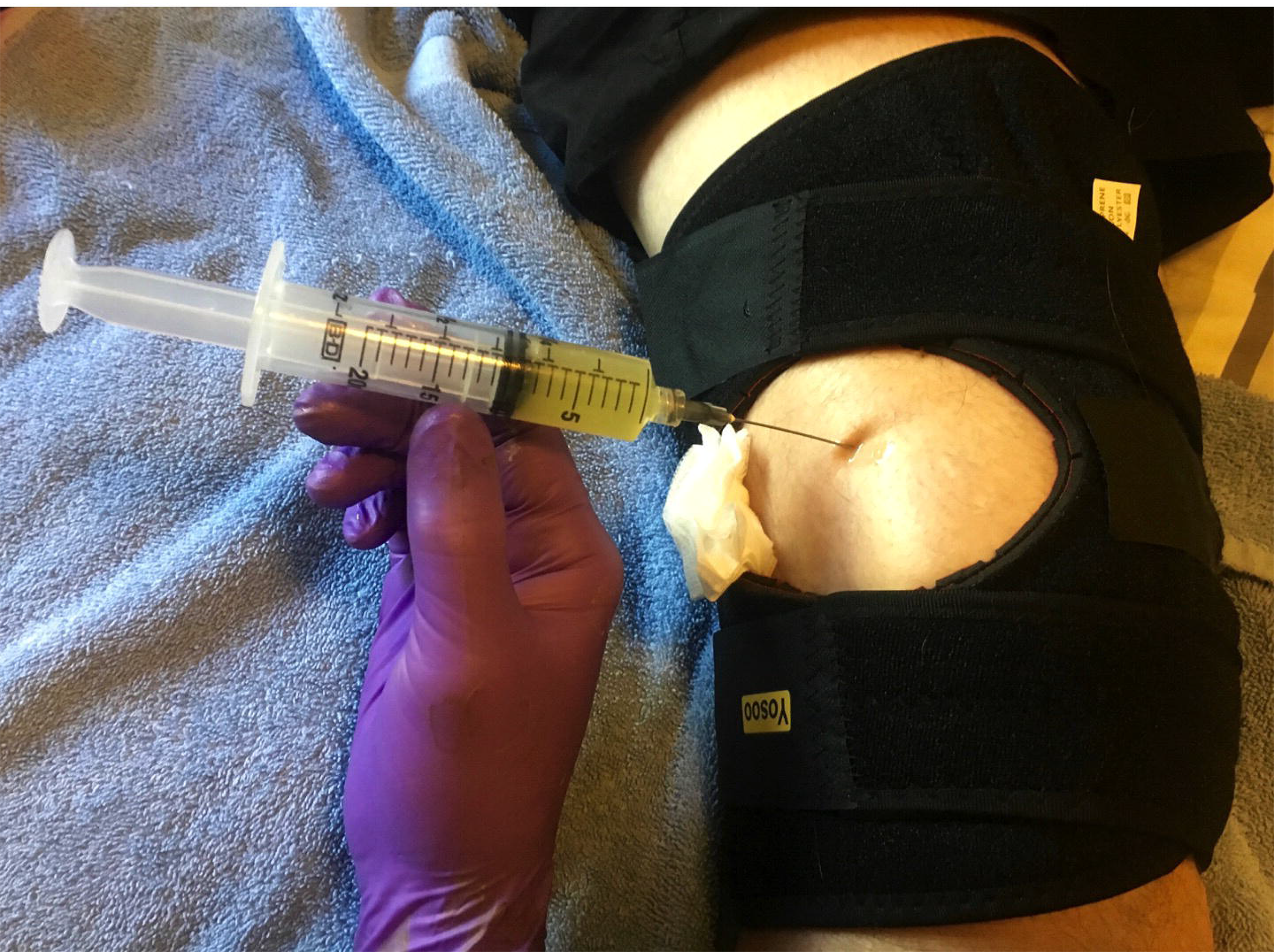
The arthrocentesis is performed through the superiolateral approach with an elastomeric brace applying circumferential (radial) constant mechanical compression to the knee and forcing residual fluid where it can be accessed at the superiolateral portal. An absorbent impermeable drape can be placed in the access portal to protect the brace.

**FIGURE 2.**
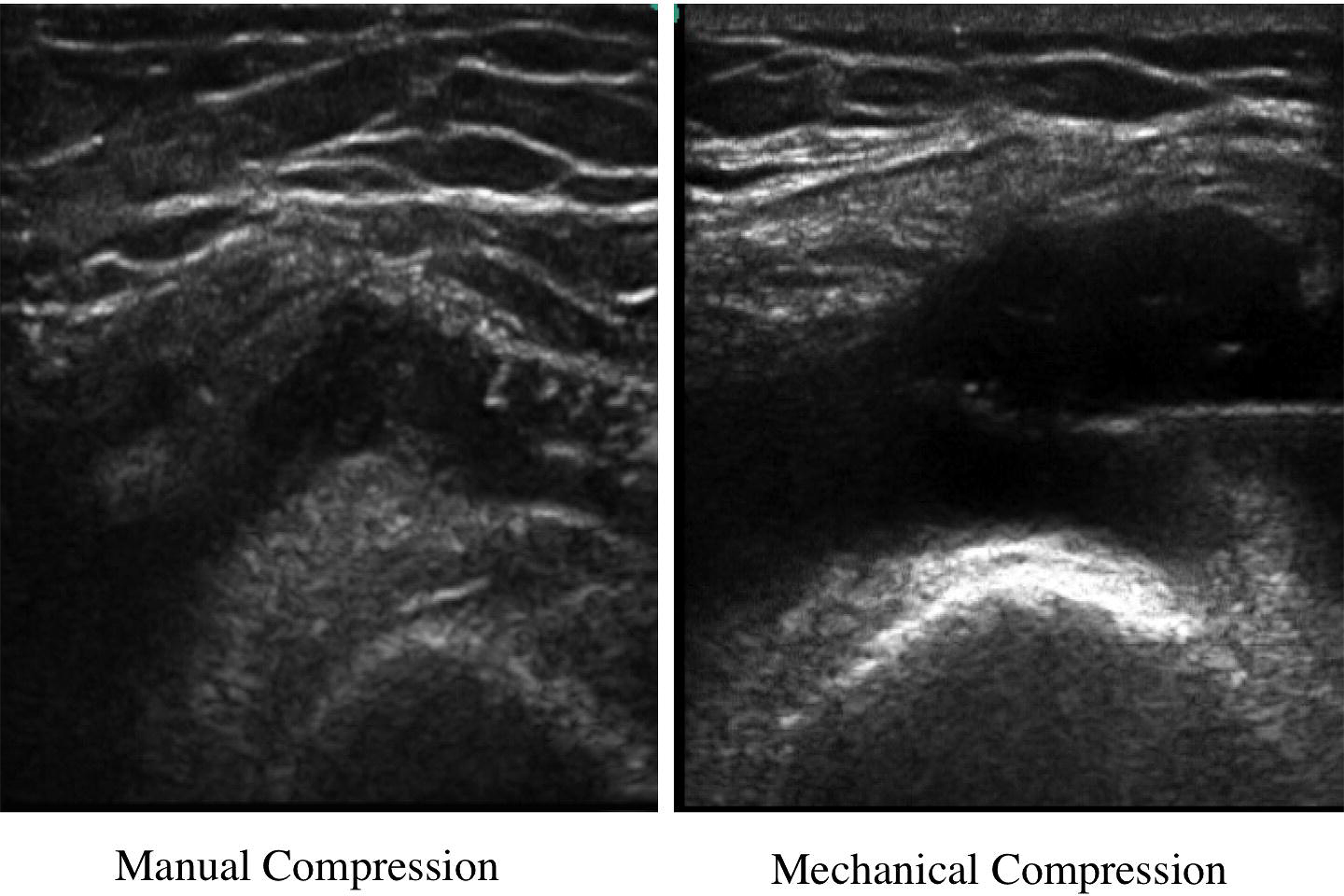
This ultrasound image demonstrates the lateral recess of the suprapatellar bursa with manual compression versus mechanical constant compression with the elastomeric brace showing substantial shift of intraarticular synovial fluid toward the access point at the lateral recess of the suprapatellar bursa. The aspiration/injection needle can be seen in the effusion on the right hand side after the constant compression brace has been applied.

### Outcome Measures

Patients were observed for serious adverse events. Diagnostic fluid was defined as ≥ to 2.0 milliliters (ml) (1 ml for culture, and 1 ml for cell counts and crystal examination [8]. Patient pain was measured with the 0-10 cm Visual Analogue Pain Scale (VAS Pain Scale), where 0 cm = no pain and 10 cm = unbearable pain [18-20]. The total number of corticosteroid injections per year were recorded in each individual including those administered in different clinics leveraging the electronic medical record to tract comprehensive quality of care as described by Schmajuk and colleagues [21]. Weitoft et al has demonstrated that complete arthrocentesis extends the time-to-next-symptomatic-flare and thus the time-to-next-intervention [5]. Time-to-next-intervention also corresponds to the time to the next corticosteroid or hyaluronan injection, surgical intervention, physical therapy or splint referral, joint imaging, or referral to another specialist, all of which increase costs. Thus the time-to-next-intervention permits preliminary baseline cost determinations [5,18,22-24].

### Costs associated with the Quality Intervention

Costs of the procedure in US dollars ($) were defined as those costs reimbursed by 2017 Medicare (United States) national rates for HCPC/CPT 20610 code for a large joint arthrocentesis for a physician office ($68.97/procedure), 15 minute outpatient encounter ($76.69), 60 mg triamcinolone acetonide ($13.00/procedure), and compressive brace ($10.00/brace) [25]. Yearly costs were calculated by multiplying the costs/procedure x 12 months divided by the months to-the-next-intervention (time to reinjection, surgical intervention, physical therapy or splint referral, joint imaging, or referral to another specialist) – this calculation is an actual underestimate of true medical costs but provides a standardization of the lowest level of costs from an injection intervention as previously described [5,20,24,25].

#### Statistical analysis

Data were entered into Excel (Version 5, Microsoft, Seattle, WA), and analyzed in SISA (Simple Interactive Statistical Analysis, http://www.quantitativeskills.com/sisa/). A power calculation was made using preliminary data at this level where α=0.0001, power = 0.9, and allocation ratio = 1.0 indicated that n=100 in each group would provide statistical power at the p<0.001 level and n = 200 in each group at the p<0.0001 level. Pierson Chi square two by two table analysis was performed on categorical data calculating both p values and confidence intervals with significance reported at the P <0.05 level. Measurement data was analyzed using the t-test calculating both p values and confidence intervals.

## RESULTS

The two study cohorts (215 conventional and 215 compression-assisted) were similar in demographics and baseline pain measures. The mean age of the conventional cohort was 63.2±11.6 years and the compression-assisted cohort was 60.1±12.1 years (p=0.007, 95% CI of difference -5.3 <-3.1< −0.9). Male:female ratio was 17:198 (92% female) in the conventional cohort, and 19:196 (88% female) in the compression cohort (p=0.73, z for 95% CI= 1.96, Pierson), typical of studies of osteoarthritis of the knee demonstrating a female gender bias. Pre-procedural pain according to the 10 cm VAS was 7.2±1.9 cm in the conventional cohort and 7.2±1.8 cm in the compression cohort (p =0.87, 95% CI of difference:−0.4 <0.0< 0.3 (Wald)). Procedural pain was 2.6±1.6 cm in the conventional cohort and 2.4±1.8 cm in the compression cohort (p = 0.22, 95% CI of difference −0.5 < −0.2< 0.1 (Wald)). Post-Procedural pain was 1.1±1.5 cm in the conventional cohort and 1.0±1.6 cm, in the compression cohort (p = 0.5, 95% CI of difference −0.4 < -0.1< 0.2 (Wald)).

There were no serious adverse events encountered by the 430 patients in the cohort including but not limited to reaction to local anesthesia, needle-stick, infection, septic joint, hemarthrosis, deep venous thrombosis, pseudoseptic arthritis, dermal atrophy, significant bruising, hemorrhage or post-injection visits to emergency facilities.

Diagnostic synovial fluid (≥2 ml) was obtained in 9.3% (20/215) without compression and 40.9% (88/215) with compression (p=0.00001, z for 95% CI= 1.96, Pierson). Absolute volume of arthrocentesis fluid yield without compression was 1.6±6.4 ml versus 5.3±11.2 ml with compression (231% increase, CI of difference 2.0 <3.7< 5.4, p=0.00001). These data are shown in Figure 3 in graphic form.

**FIGURE 3.**
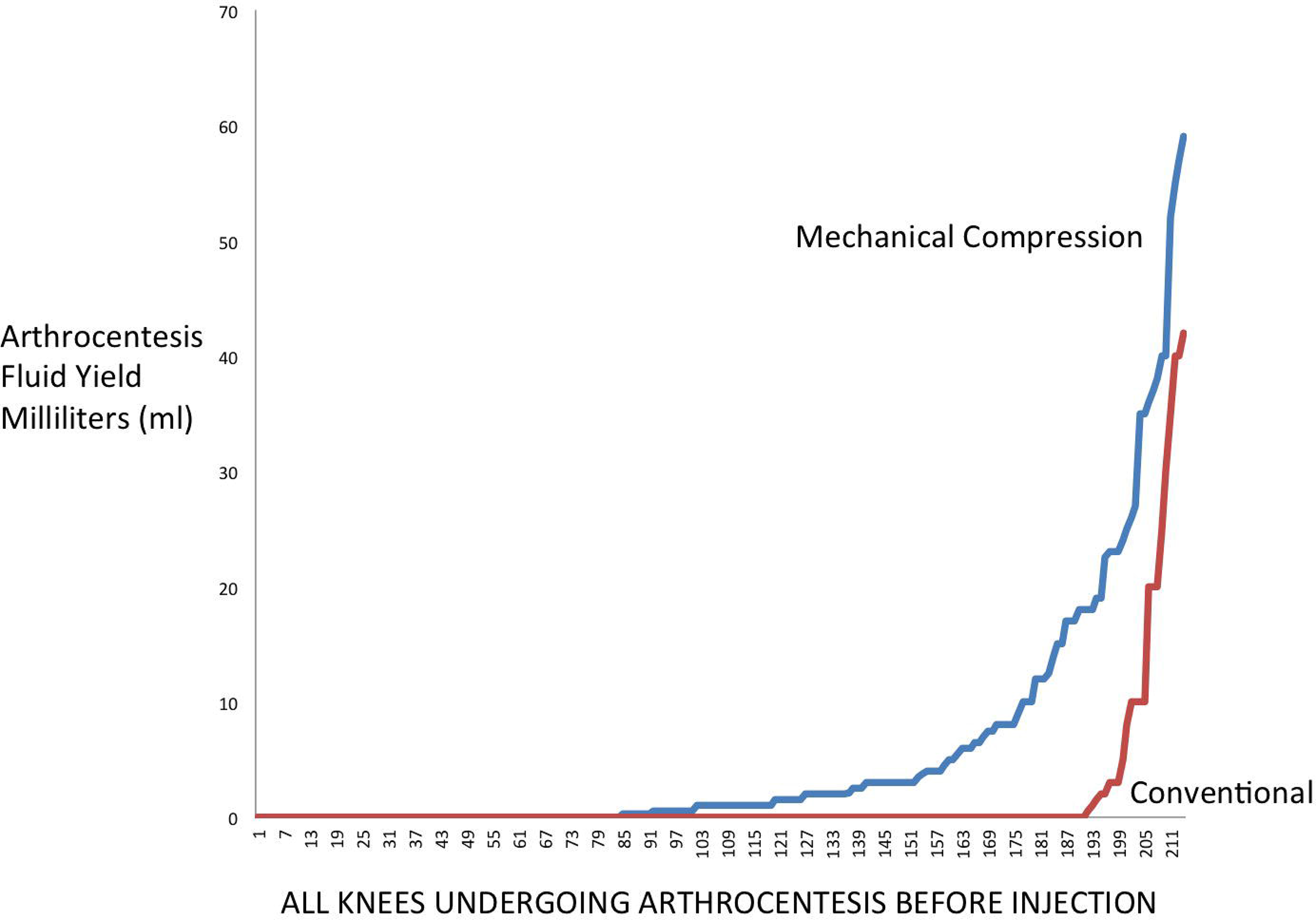
This graph shows the markedly increased synovial fluid yield with mechanical constant compression (upper line) (n=215) and conventional (lower line) (n=215), demonstrating a mean 230% increase in synovial fluid yield (p<0.0001).

In the palpably non-effusive (dry) knees (n =177) the absolute volume of arthrocentesis fluid obtained without compression was 0.0±0.00 ml versus 1.5±2.4 ml with compression (n = 178) (>100 % increase, 95% CI of difference 0.995 <1.5< 2.005, p=0.0001).

In the palpably effusive knee absolute volume of arthrocentesis without compression (n=38) was 14.7±13.8 ml versus 25.3±15.5 ml with compression (n = 37) (72.1% increase, 95% CI of mean difference: −3.0 <-1.7< −0.3, p=0.02).

Mechanical compression was associated with fewer subsequent corticosteroid injections per year: mechanical compression: 1.7±0.9 injections/year as opposed to conventional: 2.4±0.5 injections/year (p<0.00001, 95% CI of difference −0.83<−0.70<−0.56).

Time-to-next-intervention after intraarticular corticosteroid injection was longer with constant compression 6.9±3.5 months as opposed to conventional treatment 5.1±2.7 months (p<0.00001, 95% CI of difference 1.2<1.8< 2.3). These data are shown in Figure 4 in graphic form.

**FIGURE 4.**
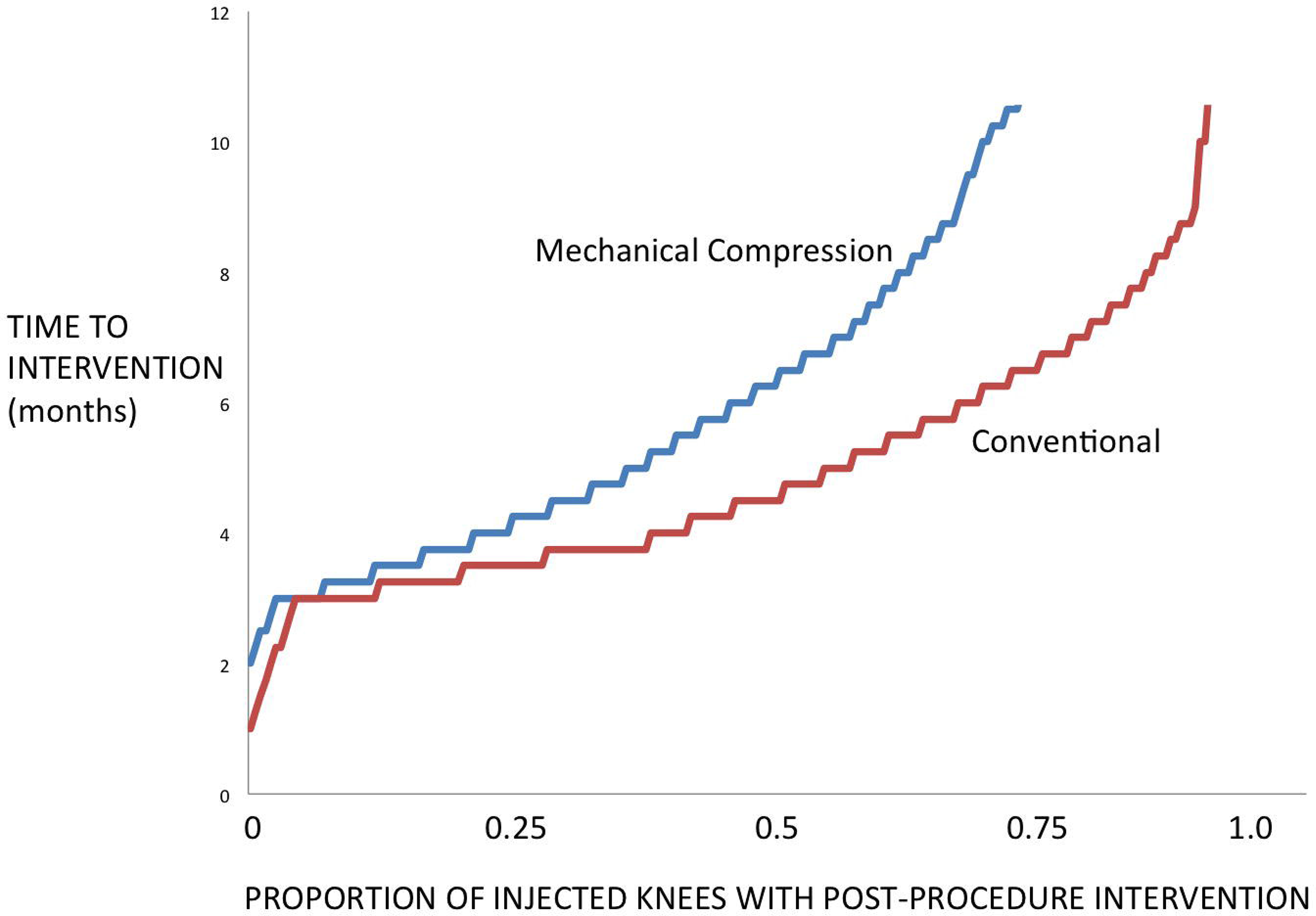
This graph shows the increased time-to-next intervention with mechanical constant compression (upper line) (n=215) and conventional (lower line) (n=215) in 430 knees demonstrating a mean 35% increase in time-to-next-intervention (p<0.00001).

Time-to-next-intervention was also longer in the effusive knees treated with mechanical compression, 7.3±3.0 months as opposed to conventional 5.6±3.0 months (p<0.016, 95% CI of difference-3.1 <-1.7< -0.3).

Time-to-next-intervention was also longer in the 178 non-effusive knees treated with constant compression, 6.9±3.5 months as opposed to 177 non-effusive knees who receive conventional treatment 5.0±2.6 months (p<0.01, 95% CI of difference -3.3 <-1.9< -0.5).

Lowest level costs per year for conventional knee aspiration/injection was $373±189/year/knee; and for mechanical compression was $293±152/year/knee (p<0.0001, 95% CI of difference 47 <80< 112), thus, mechanical compression did not increase yearly costs.

## DISCUSSION

In an effort to improve the arthrocentesis process, Meehan et al have recently demonstrated that an external compressive brace shifts fluid in the knee where it can be more easily accessed [7]. Similarly, Bhavsar et al demonstrated that external mechanical compression of the knee collapsed non-target synovial spaces and dilated and expanded target spaces and thus improved arthrocentesis success and yield in both the effusive and non-effusive knee [8]. Yaqub et al have also demonstrated that mechanical compression permits improved arthrocentesis success with the knee in different positioning [9]. Because of low cost and improved arthrocentesis success it has been proposed that mechanical compression could be used as a low cost quality improvement intervention in procedure clinics [7-9].

As part of an institutional quality improvement program to reduce occupational needle-sticks and optimize needle procedures of the knee, we introduced mechanical compression of the osteoarthritic knee before arthrocentesis and corticosteroid injection. Meehan et al, Bhavsar et al, and Yaqub et al used a repeated measure paired study design in the same subject that was sensitive to aspiration success, but not to other outcomes, costs, or changes in adverse events [7-9]. The present pragmatic study used a different design to study the quality effects of mechanical compression, specifically a two-cohort design typical of quality improvement interventions [7-10,21-26].

In the present two-cohort study mechanical compression was associated with an increased diagnostic joint aspiration success from 9% to 44%, and the volume of fluid collected by 231% (Figure 3), similar to the findings of Bhavsar et al who used a repeated measure in the same individual study design [8]. The use of the mechanical compression brace did not appear to have a negative effect on subsequent injection outcomes. Time-to-next-intervention after compression-assisted arthrocentesis was 6.9±3.5 months compared to conventional arthrocentesis: 5.1±2.7 months (p<0.00001). Mechanical compression was associated with in a reduction in the total number of corticosteroid injections administered per year: mechanical compression: 1.7±0.9 injections/year; conventional: 2.4±0.5 injections/year (p<0.00001). Mechanical compression did not increase overall costs associated with management of the knee (mechanical compression: $293.30/year/knee, conventional: $373.29/year/knee, p<0.0001). Thus, mechanical compression of the knee for arthrocentesis before injection appears to be a reasonable quality improvement intervention for knee procedures that improves quality measures and does not increase overall costs.

A number of prior studies have focused on the need to accurately place the needle intraarticularly and completely aspirate as much effusion as possible prior to injection, and all have generally demonstrated improved joint injection outcomes [5,18,27-29]. In 2003 Weitoft et al demonstrated that complete aspiration of the knee prior to corticosteroid injection prolonged the time-to-flare, and thus, reduced the need for repetitive corticosteroid injection [5]. If the data in Figure 4 of the present study are directly compared to the published data in Figure 1 of Weitoft et al, the present independent data show a similar relationship in regards to improvement in outcome after complete aspiration [5].

Extraction of joint fluid for biomarker analysis even in the dry (non-effusive) knee is presently an important area in arthritis research and may also be integral to future joint preservation strategies and therapies [6,22,29,30]. Similar to Bhavsar et al, Yaqub et al, and Meehan et al, but using a different study design, we found that mechanical compression permitted fluid extraction in a much greater proportion of osteoarthritic knees that presented clinically as dry, non-effusive knees (Figure 3) [7-9]. The failure of full synovial fluid extraction during conventional aspiration of both the effusive and non-effusive knee may be due to a combination of mistargeting of the needle, the complex intraarticular synovial compartments that trap viscous fluid, ineffective manual compression, resistance to movement of fluid due to the semi-solid gel-like properties of synovial fluid, and its non-Newtonian elasticity and viscosity [8,29-31]. A constant circumferential pressure as provided by a mechanical compressive brace dilates the joint space to be targeted (Figures 1 and 2) and takes advantage of the classic rheological properties of synovial fluid to permit linear non-turbulent fluid flow to the needle access point, enhancing fluid aspiration (Figure 3).

One limitation of this study is that medial approaches to aspiration and injection were not used; it is possible that medial approaches might be just as successful by placing the compressive brace in a reverse orientation. Another potential limitation was the sequential rather than randomized study design that potentially could be a cause of consistent bias; however, sequential methodology is a standard quality improvement vehicle in all hospitals [10,22,23,26]. In the present study, an elastomeric brace was used, it is anticipated that commercially available pneumatic braces would have similar mechanical and clinical effects [32].

## Conclusion

Mechanical compression of the knee prior to corticosteroid injection permits more complete synovial fluid extraction, and does not appear to increase costs or adversely affect outcomes. Routine mechanical compression of the knee for arthrocentesis and injection is a reasonable quality improvement intervention in a procedure clinic.

### Funding

There was no funding for this study.

